# The biosynthetic origin of psychoactive kavalactones in kava

**DOI:** 10.1101/294439

**Authors:** Tomáš Pluskal, Michael P. Torrens-Spence, Timothy R. Fallon, Andrea De Abreu, Cindy H. Shi, Jing-Ke Weng

## Abstract

For millennia, humans have used plants for medicinal purposes. However, our limited understanding of plant biochemistry hinders the translation of such ancient wisdom into modern pharmaceuticals^1^. Kava (*Piper methysticum*) is a medicinal plant native to the Polynesian islands with anxiolytic and analgesic properties supported by over 3,000 years of traditional use as well as numerous recent clinical trials^2–5^. The main psychoactive principles of kava, kavalactones, are a unique class of polyketide natural products known to interact with central nervous system through mechanisms distinct from those of the prescription psychiatric drugs benzodiazepines and opioids^6,7^. Here we report *de novo* elucidation of the biosynthetic pathway of kavalactones, consisting of seven specialized metabolic enzymes. Based on phylogenetic and crystallographic analyses, we highlight the emergence of two paralogous styrylpyrone synthases, both of which have neofunctionalized from an ancestral chalcone synthase to catalyze the formation of the kavalactone scaffold. Structurally diverse kavalactones are then biosynthesized by subsequent regio- and stereo-specific tailoring enzymes. We demonstrate the feasibility of engineering heterologous production of kavalactones and their derivatives in bacterial, yeast, and plant hosts, thus opening an avenue towards the development of new psychiatric therapeutics for anxiety disorders, which affect over 260 million people globally^8^.

## Main text

To investigate kavalactone biosynthesis, we first surveyed the occurrence of kavalactones in kava (Figure 1A) and four related *Piper* species by liquid-chromatography mass-spectrometry (LC-MS). While kavalactones are among the most abundant metabolites found in kava, they are absent in the other examined species (Figure 1B), suggesting their underlying biosynthetic machinery likely emerged after the diversification of the *Piper* genus. We also detected flavokavain A, B, and C in kava, which are specialized chalconoids with reported anti-cancer properties^9,10^ (Figure 1B).

**Figure 1.**
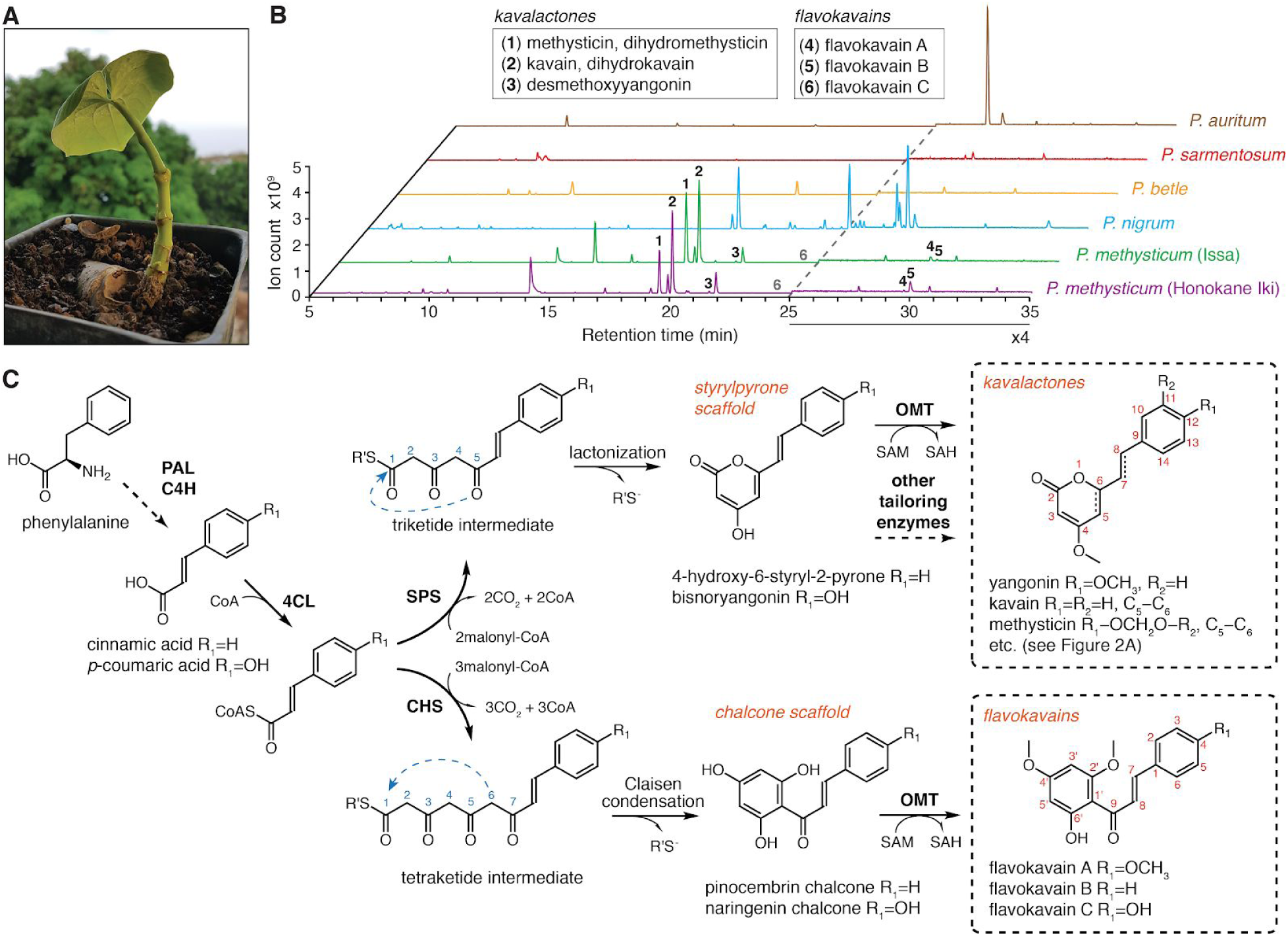
Chemotypes of select *Piper* species and the bifurcation of kavalactone and flavokavain biosynthesis from common hydroxycinnamoyl-CoA precursors in kava. **A**. A young root cutting of the Hawaiian kava cultivar *Honokane Iki*. **B.** LC-MS base peak plots of leaf tissue extracts from two different cultivars of kava (*Piper methysticum*) and four other *Piper* species showing that kavalactones and flavokavains are major secondary metabolites in kava absent in the other examined *Piper* species. The intensity in the 25-35 min region was increased 4-fold to help visualize the flavokavain peaks. The flavokavain C (**6**) peak is present but below the plot baseline. **C.** Hypothesized mechanisms for kavalactone and flavokavain biosynthesis. The polyketide cyclization mechanisms are denoted by blue arrows. PAL, phenylalanine ammonia-lyase; C4H, *trans*-cinnamate 4-monooxygenase; 4CL, 4-coumarate-CoA ligase; SPS, styrylpyrone synthase; CHS, chalcone synthase; OMT, *O*-methyltransferase; CoA, coenzyme A; SAM, *S*-adenosyl-L-methionine; SAH, *S*-adenosyl-L-homocysteine.

Considering the structural relationship between kavalactones and flavokavains, which feature the styrylpyrone and chalcone backbones, respectively, we hypothesized that kavalactone biosynthesis likely involves styrylpyrone synthase (SPS), a polyketide synthase related to chalcone synthase (CHS), which catalyzes the first committed enzymatic step in flavonoid biosynthesis and is ubiquitously present in all land plants^11^(Figure 1C). Like CHS, SPS would accept a 4-coumarate-CoA ligase (4CL)-generated hydroxycinnamoyl-CoA thioester (e.g., *p*-coumaroyl-CoA) as a starter substrate, and catalyzes iterative decarboxylative condensations of two-carbon ketone units derived from its co-substrate, malonyl-CoA. While CHS produces a tetraketide intermediate that further cyclizes via Claisen condensation to yield the chalcone backbone, SPS would yield a triketide intermediate that lactonizes to form the styrylpyrone backbone (Figure 1C). Additional tailoring enzymes, such as *O*-methyltransferases (OMTs) and oxidoreductases, are necessary to further decorate the initial scaffolds to give rise to the full repertoire of kavalactones and flavokavains present in kava plants (Figure 2).

**Figure 2.**
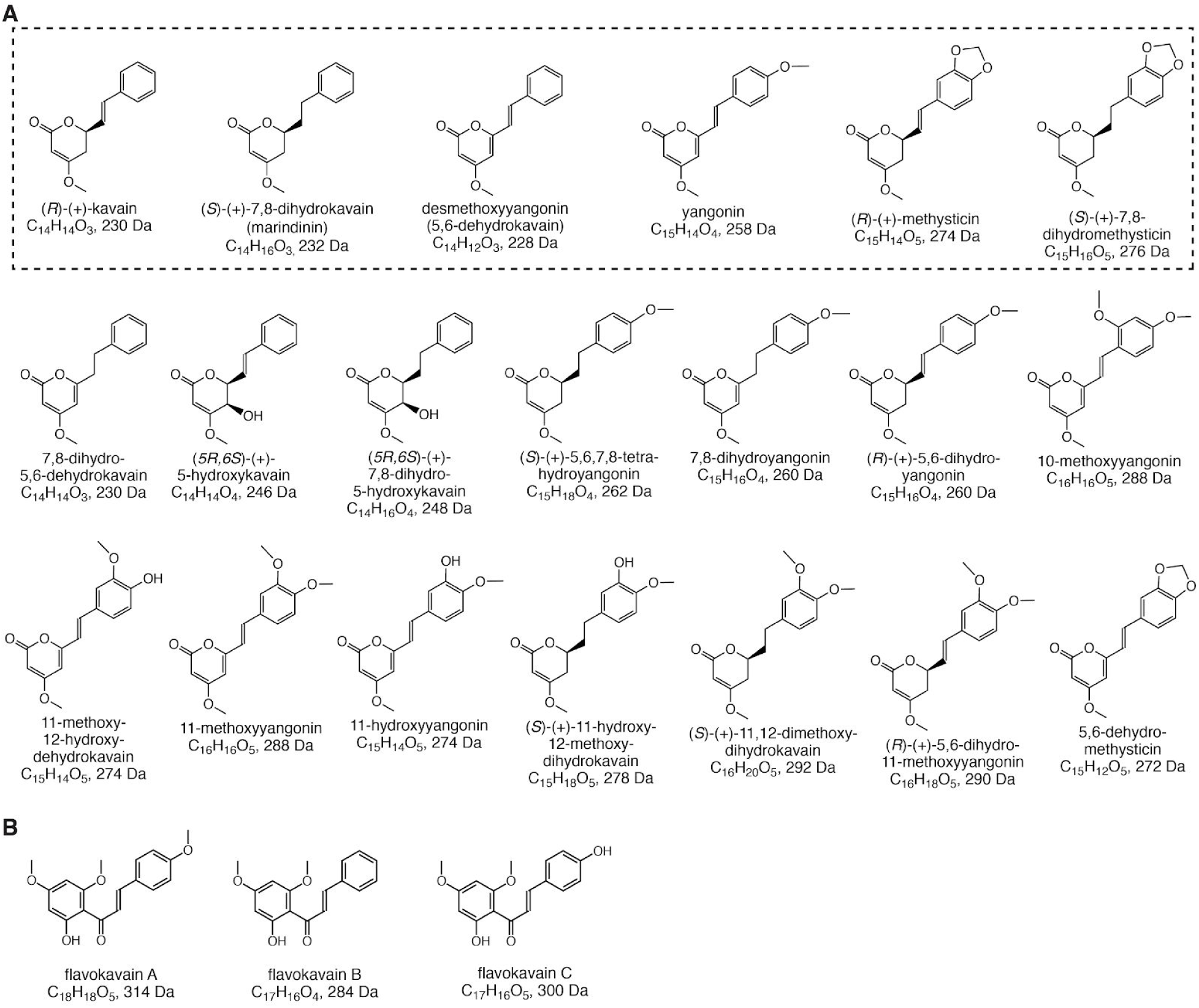
The chemical diversity of kavalactones and flavokavains. **A.** Chemical structures, formulas, and molecular masses of twenty known kavalactones. The six major kavalactones, which constitute over 96% kavalactone content in the plant rhizome, are highlighted within a dashed rectangle. **B.** Chemical structures, formulas, and molecular masses of three known flavokavains.

Kava is a decaploid (2n = 10x = 130 chromosomes)^12^, rendering biosynthetic inquiry via conventional genetic approaches infeasible. We therefore turned to a multi-omics-guided candidate gene approach^13–15^. We first identified three *CHS*-like genes in a kava transcriptome assembled *de novo* from leaf and root tissues (Supplementary Table 1). Phylogenetic analysis of these genes in the context of other land plant *CHS*s suggests that two of the three genes were likely derived from recent gene duplication events specific to kava (Figure 3A). *In vitro* enzyme assays using heterologously expressed and purified enzymes established that these two *CHS*-like genes indeed encode functional SPSs. Both enzymes exclusively produce the triketide lactone bisnoryangonin from *p*-coumaroyl-CoA (Figure 3B), and are therefore named *Pm*SPS1 and *Pm*SPS2 hereafter. The assay also established that the third *CHS*-like gene encodes a bona fide CHS, which produces the expected tetraketide-derived naringenin chalcone, and is referred to as *Pm*CHS hereafter (Figure 3B). It is noted that *Pm*CHS also produces *p*-coumaroyltriacetic acid lactone (CTAL) and bisnoryangonin in addition to naringenin chalcone *in vitro*, although similar derailment products have been reported for a number of previously characterized CHSs^16^.

**Figure 3.**
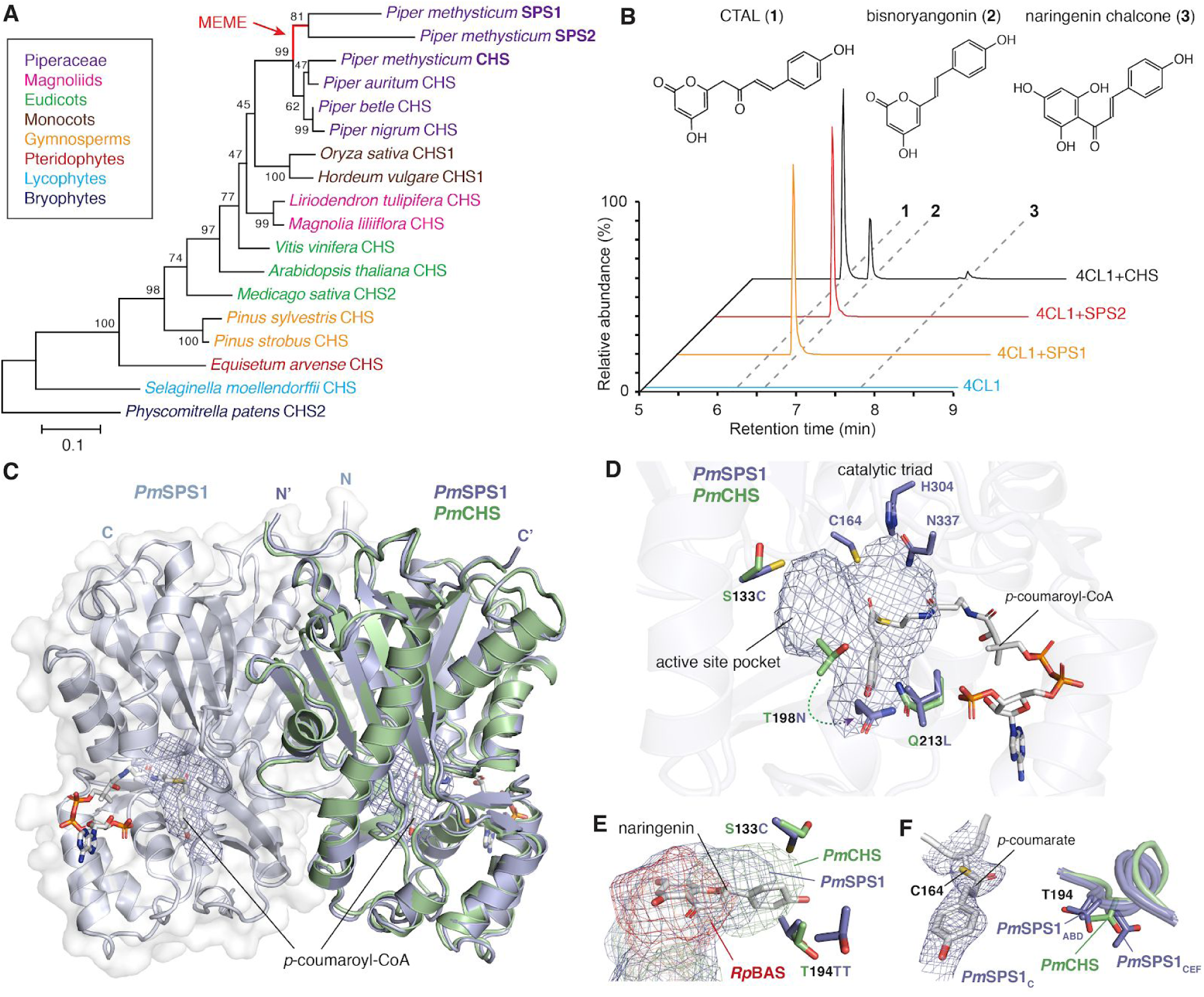
Mechanistic basis for the neofunctionalization of SPSs from ancestral CHS in kava. **A.** Maximum-likelihood phylogenetic analysis the three CHS-like sequences (*Pm*SPS1, *Pm*SPS2, and *Pm*CHS) from kava with other CHS otholog sequences from select land plant lineages. Bootstrap values (based on 1000 replicates) are indicated at the tree nodes. The branch undergone episodic positive selection as detected by the Mixed Effects Model of Evolution analysis^19^ is highlighted in red and denoted with MEME. **B.** Combined LC-MS extracted ion chromatograms (XICs) of 273.076 m/z (naringenin chalcone and CTAL) and 231.065 m/z (bisnoryangonin) showing the *in vitro* activities of *Pm*SPS1, *Pm*SPS2, and *Pm*CHS from a 4CL-coupled enzyme assay using *p*-coumaric acid as the starter substrate. **C.** An overlay of the *Pm*CHS apo structure and the *Pm*SPS1-*p*-coumaroyl-CoA holo structure (RMSD=1.019 Å). Both proteins form homodimers (only a single protomer of *Pm*CHS is shown). Active-site pocket is indicated with a mesh. **D.** Structural features of the *Pm*SPS1 active site. The three active-site amino acid substitutions between *Pm*SPS1 and *Pm*CHS are highlighted. **E.** The C133 and T194 residues of *Pm*SPS1 dictate the size and shape of the *Pm*SPS1 active site pocket in comparison to *Pm*CHS and *Rheum palmatum* BAS (*Rp*BAS). The naringenin coordinates are from the *Medicago sativa* CHS structure (PDB 1CGK^64^) based on structural alignment. **F.** *p*-Coumaroyl-monoketide intermediate covalently bound to the catalytic cysteine as captured in chain C of the *Pm*SPS1-*p*-coumaroyl-CoA structure. The |2*F*_*o*_ – *F*_*c*_*|* electron density map is contoured at 1.2 σ. T194 residue of a nearby loop captured in closed (chains A, B, and D) and open (chains C, E, and F) conformations, in comparison with the corresponding residue in the *Pm*CHS structure.

To assess the *in vivo* function of the three newly identified kava PKSs, we expressed each of them as stable transgenes in the *Arabidopsis thaliana CHS*-null mutant *tt4-2* background^17^.

While *PmCHS* restored flavonoid biosynthesis in *tt4-2*, neither *PmSPS1* nor *PmSPS2* did, consistent with their respective *in vitro* biochemical activities (Figure 4). However, we could detect neither bisnoryangonin nor other styrylpyrones in transgenic Arabidopsis expressing *PmSPS1* or *PmSPS2*. Similarly, *Agrobacterium* infiltration-mediated transient expression of *PmSPS1* or *PmSPS2* in the leaves of *Nicotiana benthamiana^18^* also failed to yield any styrylpyrone compounds. Upon careful examination, we instead identified a series of benzalacetone-type metabolites that accumulated in transgenic Arabidopsis or Nicotiana plants expressing *PmSPS1* (Supplementary Note 2). We conclude that styrylpyrones generated by heterologously expressed *PmSPS1* are rapidly turned over by unknown enzymes present in Arabidopsis or Nicotiana to yield benzalacetones as breakdown products.

**Figure 4.**
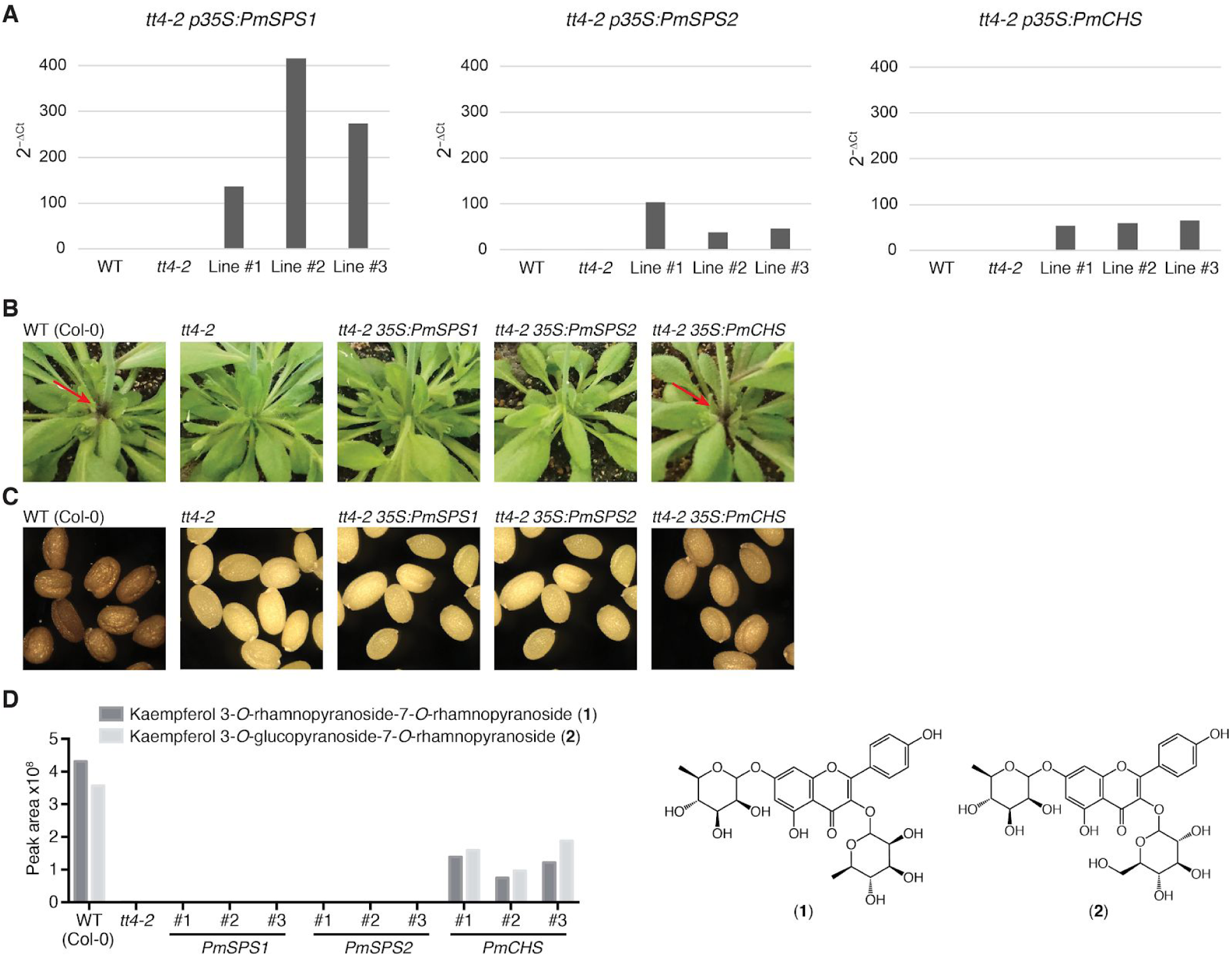
Stable transgenic expression of *PmSPS1*, *PmSPS2*, and *PmCHS* in Arabidopsis *tt4-2* background. **A.** Expression levels of *PmSPS1*, *PmSPS2*, and *PmCHS* relative to a reference gene, *At1g13320*, in three different Arabidopsis lines measured by qRT-PCR in the T2 generation. **B.** Transgenic expression of *PmCHS*, but not *PmSPS1* and *PmSPS2* rescues the anthocyanin phenotype observed at the stem base in the Arabidopsis *tt4-2* mutant background. **C.** Transgenic expression of *PmCHS* partially rescues the transparent testa phenotype of the seed coat in the Arabidopsis *tt4-2* mutant background. The difference in coloring intensity in WT and *tt4-2*-35S:*PmCHS* seed coat an be attributed to the strength of the 35S promoter in seeds. **D.** LC-MS quantification of two major flavonoids^65^ in the leaves of three independent transgenic lines of *tt4-2*-35S:*PmCHS* plants.

To probe the mechanistic basis for SPS neofunctionalization from the ancestral CHS, we solved the apo structure of *Pm*CHS and the holo structure of *Pm*SPS1 in complex with *p*-coumaroyl-CoA by X-ray crystallography (Table 1). Both proteins are homodimers and share the canonical αβαβα thiolase fold typical for plant type III PKSs^11^ (Figure 3C). An evolutionary analysis using the Mixed Effects Model of Evolution (MEME)^19^ on the ancestral branch of kava SPSs (Figure 3A) detected six amino acid residues under episodic diversifying selection (p<0.05) (Figure 5). Three of these substitutions, S133C, T198N, and Q213L, are mapped to the enzyme active site (Figure 3D). Notably, the T198N substitution was previously reported as one of the three mutations sufficient to convert CHS into a 2-pyrone synthase (2PS)^20^. We also observed an unusual insertion of threonine (T194) in a loop region abutting the active site, together with the S133C substitution, causing a reduction of the *Pm*SPS1 active site volume compared to *Pm*CHS in the direction along the polyketide chain elongation (Figure 3E). In comparison, the previously reported *Rheum palmatum* benzalacetone synthase (*Rp*BAS), which catalyzes only a single polyketide elongation^21^, features a further shortened active site compared to *Pm*SPS1 (Figure 3E). These observations therefore support the notion that subtle changes of active site volume and shape of type III PKSs dictate the iterative cycles of polyketide elongation and alternative cyclization mechanisms^22^. In addition, we observed two different conformations of T194 among the six protomers present in the asymmetric unit, suggesting the dynamic nature of the underlying loop and its potential role in the catalytic cycle of *Pm*SPS1 (Figure 3F). One of the protomers in our *Pm*SPS1-*p*-coumaroyl-CoA complex structure also captures a *p*-coumaroyl-monoketide intermediate covalently bound to the catalytic cysteine (Figure 3F), providing a rare glimpse of the active site configuration after the starter substrate loading step of the PKS catalytic cycle^22^. Homology modeling and sequence analysis of *Pm*SPS2 suggest that its active site contains a similar set of amino acid substitutions as observed in *Pm*SPS1, except for the T194 insertion (Figure 5 and 6).

**Table 1.** Crystallography data collection and refinement statistics. Statistics for the highest-resolution shell are shown in parentheses.

|  | *PmSPS1* | *PmCHS* |
| --- | --- | --- |
| **Wavelength** | 0.9793 Å | 0.9793 Å |
| **Resolution range** | 82.2 - 2.7 (2.797 - 2.7) | 62.7 - 2.91 (3.014 - 2.91) |
| **Space group** | P 31 | P 21 21 21 |
| **Unit cell** | 164.402 164.402 87.1221 90 90 120 | 50.9096 120.57 125.408 90 90 90 |
| **Total reflections** | 3027278 (290476) | 1083656 (107234) |
| **Unique reflections** | 72283 (7187) | 17602 (1726) |
| **Multiplicity** | 41.9 (40.2) | 61.6 (62.1) |
| **Completeness (%)** | 99.59 (99.31) | 99.95 (100.00) |
| **Mean I/sigma(I)** | 6.90 (1.30) | 12.54 (2.85) |
| **Wilson B-factor** |  | 44.73 |
| **R-merge** | 1.406 (4.383) | 0.926 (2.025) |
| **R-meas** | 1.424 (4.445) | 0.9341 (2.043) |
| **R-pim** | 0.2252 (0.7278) | 0.1204 (0.2639) |
| **CC1/2** | 0.775 (0.157) | 0.949 (0.614) |
| **CC\*** | 0.935 (0.521) | 0.987 (0.872) |
| **Reflections used in refinement** | 72027 (7187) | 17602 (1726) |
| **Reflections used for R-free** | 1987 (199) | 1771 (179) |
| **R-work** | 0.2111 (0.2718) | 0.2615 (0.3382) |
| **R-free** | 0.2606 (0.3123) | 0.3156 (0.3688) |
| **CC(work)** | 0.668 (0.224) | 0.921 (0.694) |
| **CC(free)** | 0.403 (0.278) | 0.881 (0.694) |
| **Number of non-hydrogen atoms** | 18141 | 5880 |
| **macromolecules** | 17603 | 5846 |
| **solvent** | 135 | 34 |
| **Protein residues** | 403 | 762 |
| **RMS(bonds)** | 2293 | 0.002 |
| **RMS(angles)** | 0.004 | 0.53 |
| **Ramachandran favored (%)** | 0.62 | 94.20 |
| **Ramachandran allowed (%)** | 92.63 | 5.80 |
| **Ramachandran outliers (%)** | 6.45 | 0.00 |
| **Rotamer outliers (%)** | 0.92 | 1.59 |
| **Clashscore** | 0.05 | 24.22 |
| **Average B-factor** | 14.56 | 47.80 |
| **macromolecules** | 38.58 | 47.89 |
| **solvent** | 38.46 | 32.71 |
| **Number of TLS groups** |  | 8 |

**Figure 5.**
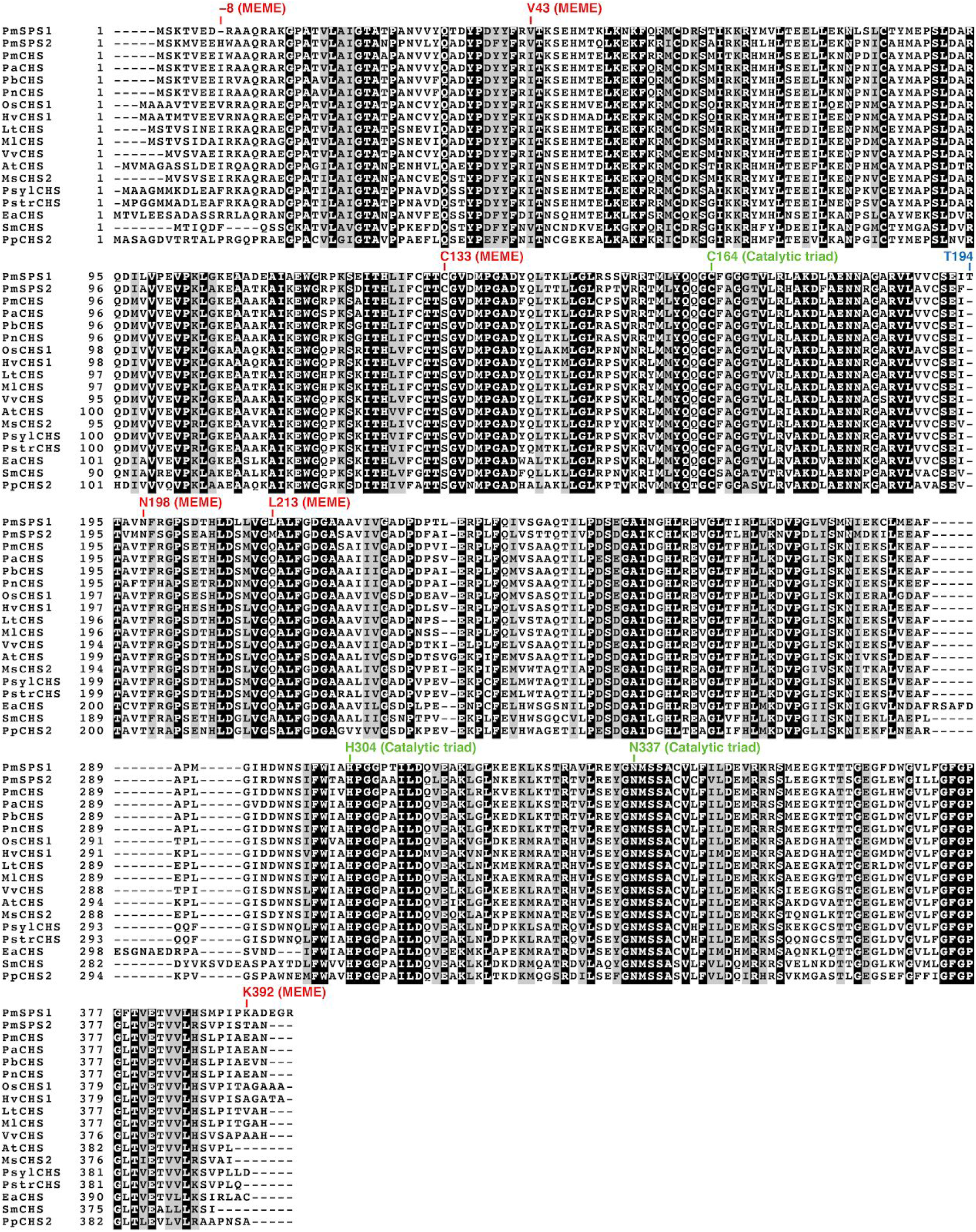
Multiple sequence alignment of CHS homologs as in Figure 3A. The residues undergone episodic positive selection were detected based on the Mixed Effects Model of Evolution^19^ along the ancestral branch of kava SPSs, and are annotated with MEME. Residue numbering is according to *Pm*SPS1 sequence.

**Figure 6.**
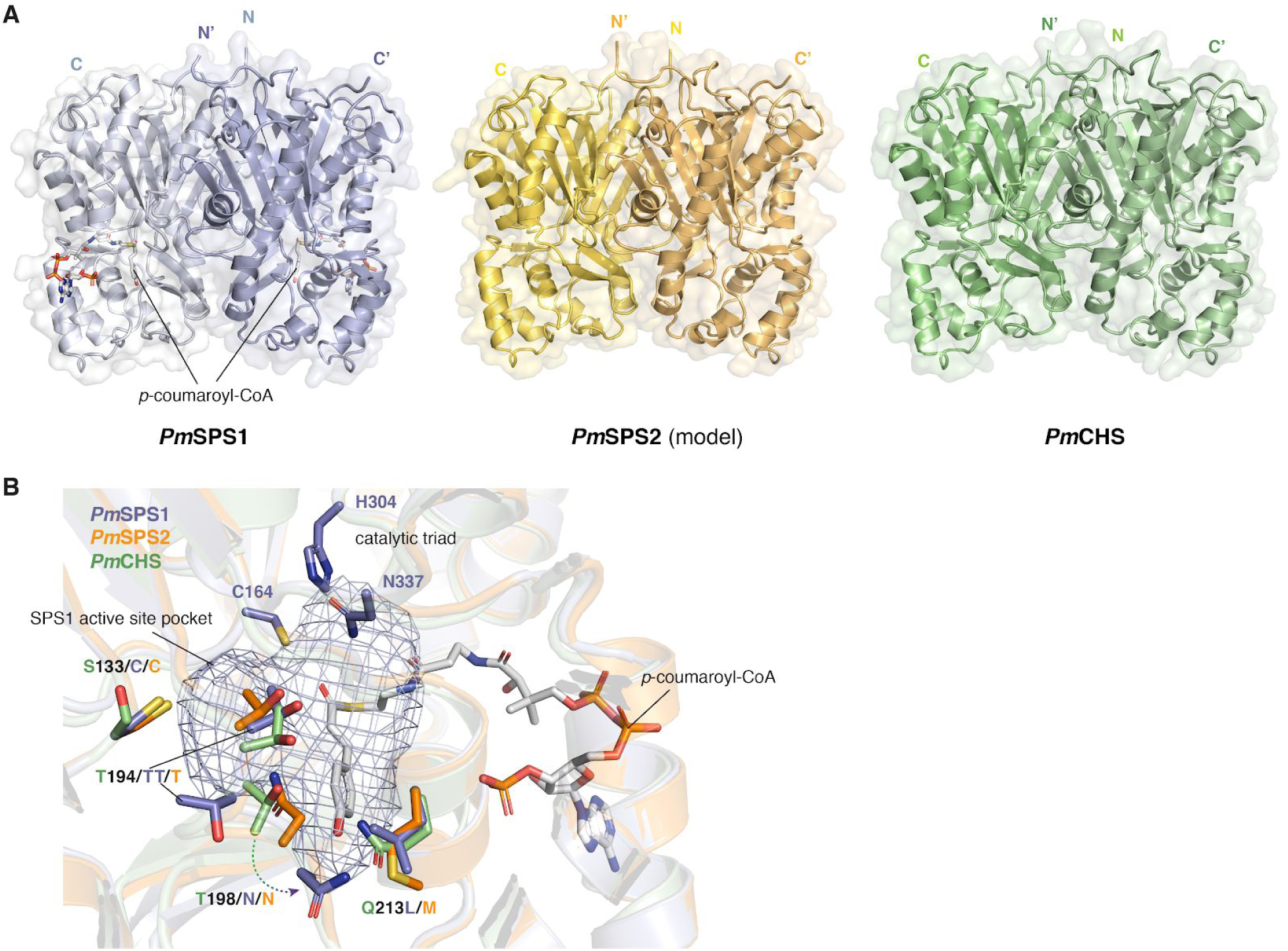
Structural comparison of *Pm*SPS1, *Pm*SPS2 and *Pm*CHS. **A.** The overall fold of experimentally determined *Pm*SPS1-*p*-coumaroyl-CoA and *Pm*CHS structures in comparison with the computationally modeled *Pm*SPS2 structure. **B.** *Pm*SPS1 and *Pm*SPS2 contain a similar set of amino acid substitutions in the active site relative to *Pm*CHS. Although both *Pm*SPS1 and *Pm*SPS2 contain the T198N substitution, the actual position of this residue is different due to the T194 insertion in *Pm*SPS1.

Structures of naturally occuring kavalactones implicate rich *O*-methylation reactions during their biosynthesis (Figure 2A). We hypothesized that kavalactone biosynthetic OMTs could have been recruited from a conserved OMT involved in plant phenylpropanoid metabolism, e.g. the caffeic acid *O*-methyltransferase (COMT) in lignin biosynthesis^23^.

Combining phylogenomics and expression analysis, we identified six putative *OMT* candidate genes from kava transcriptomes (Supplementary Table 1). Since kavalactones contain four possible *O*-methylation sites (−OH groups attached to the C_4_, C_10_, C_11_, or C_12_ positions), we established a coupled enzymatic system using a 4CL enzyme cloned from kava (designated *Pm*4CL1) and *Pm*SPS1 to produce five styrylpyrones featuring different aromatic ring modification patterns derived from five hydroxycinnamic acids, namely cinnamic, *p*-coumaric, caffeic, ferulic, and umbellic acid (Supplementary Note 3). Subsequent OMT activity assays using recombinant kava OMTs revealed that two of the six kava OMT candidates showed activity against various styrylpyrones (Figure 7A), and were thus named kava *O*-methyltransferases 1 and 2 (*Pm*KOMT1 and *Pm*KOMT2), respectively. *Pm*KOMT1 is one of the most highly expressed enzymes in kava (Supplementary Table 1), and indeed is capable of methylating the −OH groups at the C_4_, C_11_, or C_12_ positions of the styrylpyrone backbone. On the other hand, *Pm*KOMT2 activity is specific to the −OH group attached at the C_10_ position (Figure 7A). Transient co-expression of *PmKOMT1* together with *PmSPS1* or *PmSPS2* in Nicotiana resulted in the production of yangonin, a kavalactone with two methoxy groups at the C_4_ and C_12_ positions (Figure 7B), supporting its enzymatic activities observed *in vitro*. This result also suggests that *O*-methylation protects styrylpyrones produced by kava SPSs from being catabolized in Nicotiana.

**Figure 7.**
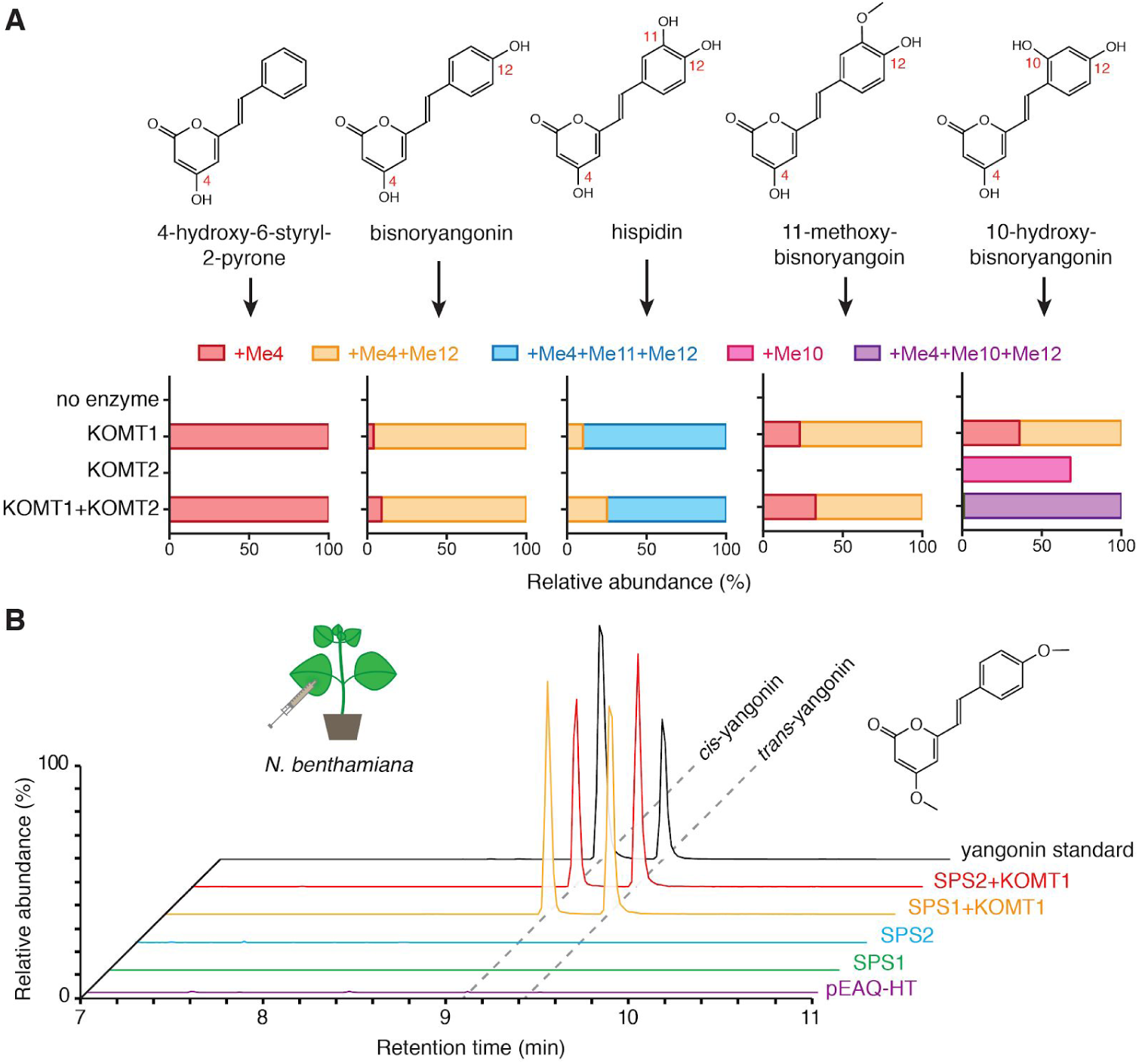
Functional characterization of kava OMTs. **A.** *In vitro* enzyme activities of *Pm*KOMT1 and *Pm*KOMT2 against five different styrylpyrone substrates. The bar charts indicate proportions of differentially methylated products relative to the unmethylated precursor (e.g., +Me4+Me12 indicates methyl group addition to both C_4_ and C_12_ sites). Detailed results are shown in Supplementary Note 3. **B.** LC-MS XICs of 259.096 m/z (yangonin) in leaf extracts from Nicotiana transiently expressing indicated enzymes. Yangonin readily isomerizes between *cis* and *trans* configurations in solution.

The styrylpyrone backbone produced by SPSs contains two olefins (C_5_=C_6_ and C_7_=C_8_), which could be reduced to alkane at either position (e.g., kavain with C_5–_C_6_ or 7,8-dihydroyangonin with C_7–_C_8_) or at both positions (e.g., 7,8-dihydrokavain) (Figure 2A). Importantly, the reduction of the 5,6-olefin occurs in a stereospecific manner, suggesting that this reaction is likely enzymatic. Using similar approaches as described above, we identified and characterized twelve kava reductases that are homologous to various known reductases in plant specialized metabolism (Supplementary Table 1). By transgenic co-expression of each of these reductase candidates together with *PmSPS1* and *PmKOMT1* in Nicotiana, we identified a single reductase candidate that produced metabolites with masses corresponding to reduced styrylpyrones (Supplementary Note 4). We thus named this enzyme kavalactone reductase 1 (*Pm*KLR1). A combined enzyme assay using purified *Pm*4CL1, *Pm*SPS1, *Pm*KOMT1, and *Pm*KLR1 enzymes starting from cinnamic acid successfully produced kavain *in vitro* (Figure 8A). Furthermore, we determined using chiral chromatography that *Pm*KLR1 produces only (*R*)-(+)-kavain, the same stereoisomer as present in kava root extract (Figure 8B). We further established that *Pm*KLR1 can only accept unmethylated styrylpyrones, and is therefore competing with *Pm*KOMT1 for the same substrates (Figure 9). This result provides an explanation to the fact that only some of the major kavalactones contain the reduced C_5−_C_6_ bond (Figure 2A).

**Figure 8.**
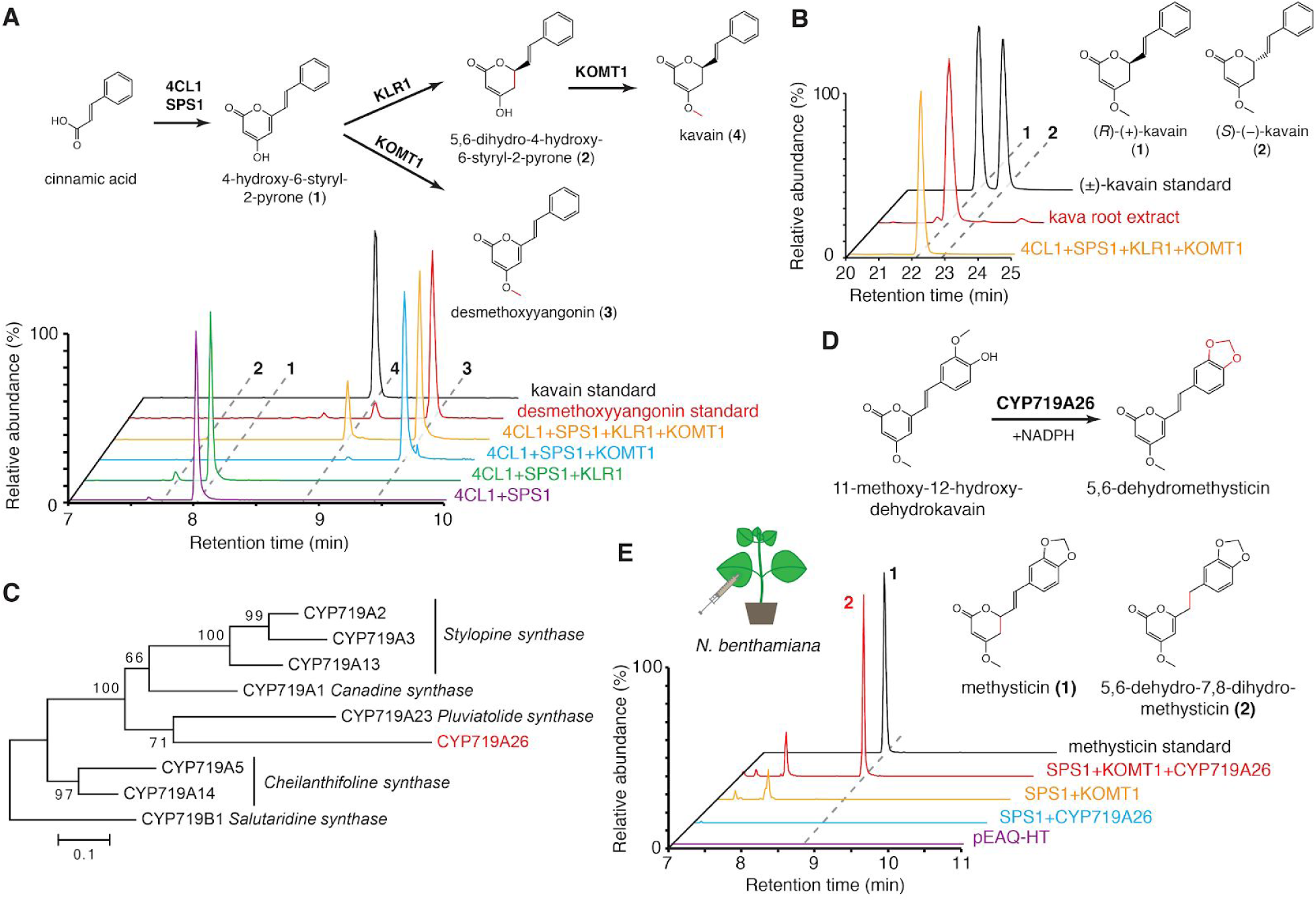
Functional characterization of kavalactone tailoring enzymes KLR1 and MTS1. **A.** Combined LC-MS XICs of 215.070 m/z (**1**), 217.086 m/z (**2**), 229.086 m/z (**3**), and 231.102 m/z (**4**) from *in vitro* enzyme assays using the indicated combinations of enzymes and cinnamic acid as a starter substrate to show the specific activity of *Pm*KLR1. **B.** Chiral chromatography-MS traces of 231 m/z (kavain) demonstrating stereo-selectivity of *Pm*KLR1. **C.** Maximum likelihood phylogenetic analysis of *Pm*CYP719A26 together with several characterized members of the CYP719 family P450s. Bootstrap values (based on 1000 replicates) are indicated at the tree nodes. The scale measures evolutionary distances in substitutions per amino acid. **D.** Proposed reaction catalyzed by *Pm*CYP719A26 (*Pm*MTS1). **E.** LC-MS XICs of 275.091 m/z (methysticin) in Nicotiana leaves infiltrated with the indicated combinations of enzymes to show the specific activity of *Pm*MTS1. The two structural isomers shown were distinguished using MS/MS analysis (Supplementary Note 5).

**Figure 9.**
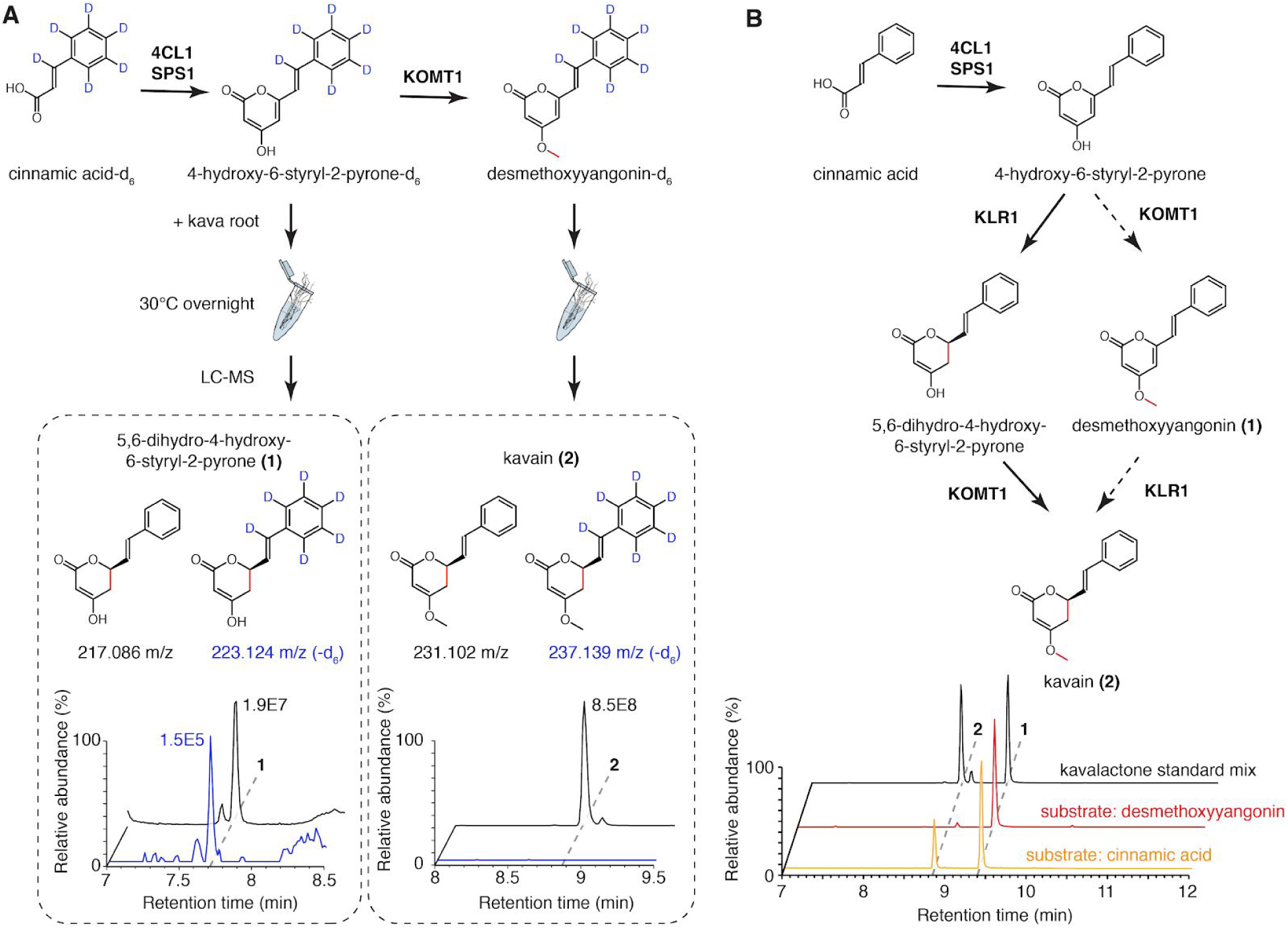
Resolving the order of *Pm*KOMT1 and *Pm*KLR1 reactions *in vivo* and *in vitro*. **A.** Isotope tracing using labeled styrylpyrone or desmethoxyyangonin precursors in live kava root suggests that only the unmethylated precursor serve as a substrate for the reduction of C_5_=C_6_ olefin. **B.** Combined LC-MS XICs of 229.086 m/z (**1**) and 231.102 m/z (**2**) from *in vitro* enzyme assays using cinnamic acid and desmethoxyyangonin as starter substrates to show that *Pm*KLR1 precedes *Pm*KOMT1. The reaction from cinnamic acid successfully produced kavain, while the same assay starting from desmethoxyyangonin failed to show any activity. The determined order of the pathway is shown with solid arrows, while the rejected order is shown with dashed arrows.

We hypothesized that the 7,8-saturated kavalactones could be derived from dihydro-hydroxycinnamoyl-CoA precursors already containing a reduced α−β bond. For example, two type III PKSs in the genera *Pinus* and *Malus* could accept dihydro-*p*-coumaroyl-CoA as a starter substrate to produce 5-hydroxylunularic acid and phloretin, respectively^24,25^. To test this, we performed coupled *Pm*4CL1−*Pm*SPS1 *in vitro* enzyme assays using phloretic acid or phenylpropanoic acid as starter substrates, both of which contain a reduced α−β bond. These assays resulted in the production of the corresponding 7,8-saturated kavalactones, respectively, suggesting that *Pm*4CL1 and *Pm*SPS1 can utilize substrates derived from dihydro-hydroxycinnamic acids with a reduced α−β bond, which are common phenylpropanoid metabolites in plants (Figure 10).

**Figure 10.**
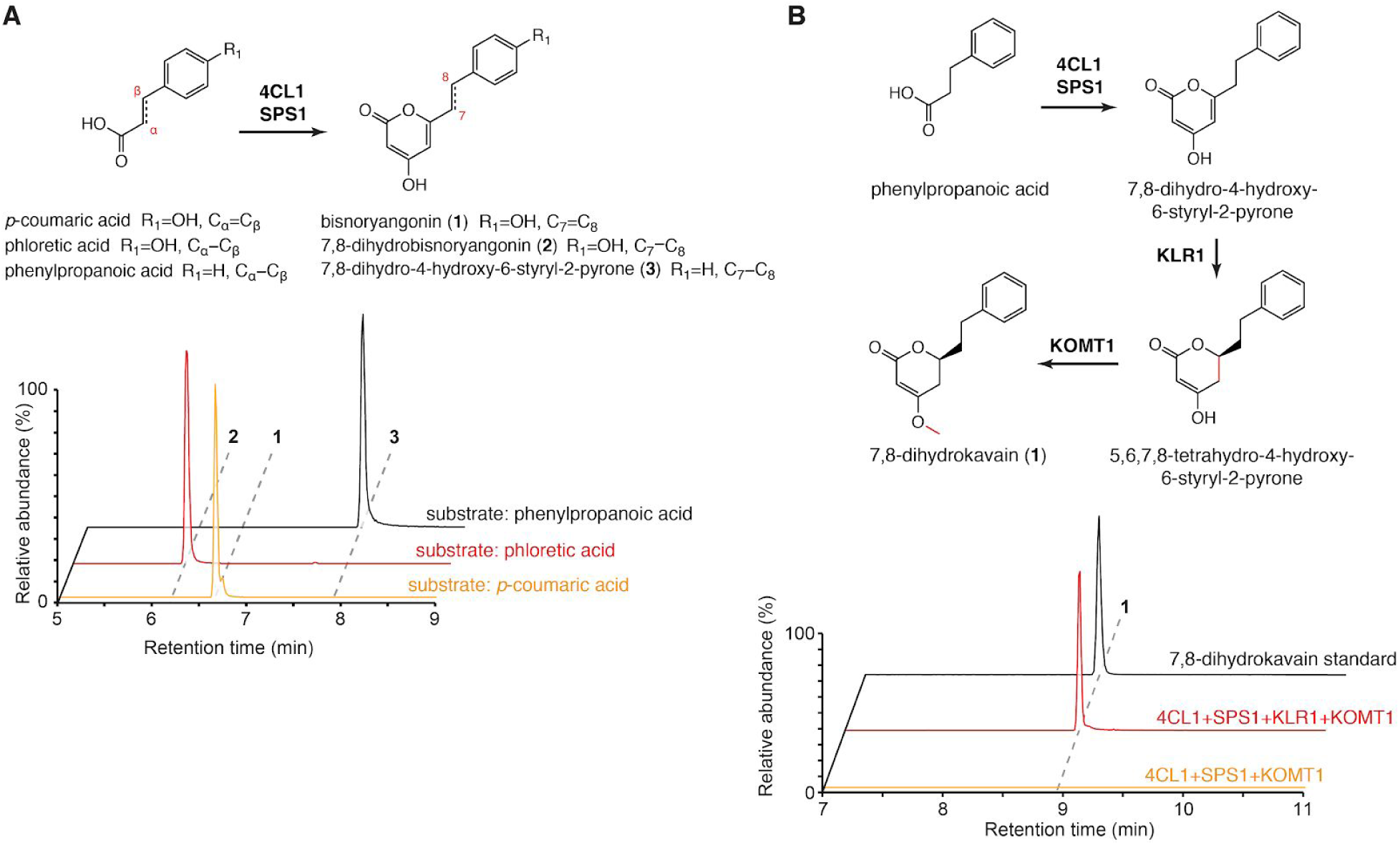
The biosynthesis of 7,8-reduced kavalactones. **A.** Combined LC-MS XICs of 231 m/z (**1**), 233 m/z (**2**), and 217 m/z (**3**) in coupled (*Pm*4CL1 + *Pm*SPS1) *in vitro* enzyme assays to show that kavalactone biosynthesis is operational starting with dihydro-hydroxycinnamic acids. The use of α,β-reduced substrates (phloretic acid and phenylpropanoic acid) resulted in the production of corresponding 7,8-reduced styrylpyrones. **B.** LC-MS XICs of 233.117 m/z (7,8-dihydrokavain) in combinatorial *in vitro* enzyme assays using the indicated enzymes and phenylpropanoic acid as a starter substrate. The addition of *Pm*KLR1 to the enzyme mix resulted in the production of 7,8-dihydrokavain, thus further supporting the conclusion that *Pm*KLR1 acts as a NADPH-dependent kavalactone 5,6-olefin reductase.

Methysticin-like kavalactones, which carry a methylenedioxy bridge at the C_11−_C_12_ position (Figure 2A), exhibit strong modulatory effects on human liver cytochromes P450, and thus contribute to the complex pharmacology of kava^26^. The CYP719 family of plant P450 enzymes is known to catalyze the formation of the methylenedioxy bridge moiety from vicinal methoxyl and hydroxyl groups, e.g. in the biosynthesis of berberine and etoposide^27,28^. We identified a single gene of the CYP719 family in kava, *PmCYP719A26* (Figure 8C), and hypothesized that it may catalyze the formation of methylenedioxy bridge using ferulic acid-derived styrylpyrone precursors carrying vicinal 11-methoxyl and 12-hydroxyl groups (Figure 8D). Indeed, co-expression of *PmSPS1*, *PmKOMT1*, and *PmCYP719A26* in Nicotiana resulted in the transgenic production of a compound with molecular mass and retention time identical to that of methysticin (Figure 8E). Fragmentation pattern comparison with a methysticin standard further identified this compound as 5,6-dehydro-7,8-dihydromethysticin (Supplementary Note 5). We therefore designated *Pm*CYP719A26 as methysticin synthase 1 (*Pm*MTS1).

Through OMT activity screen, we found that *Pm*KOMT1 and *Pm*KOMT2 also exhibit *O*-methylation activities against naringenin chalcone produced by CHS (Supplementary Note 6). Using enzyme assays and heterologous expression in Nicotiana, we confirmed that *Pm*KOMT1 methylates the −OH groups at the C_4_ and C_4’_ positions on the chalcone scaffold, while *Pm*KOMT2 methylates the −OH group at the C_2’_ position (Figure 11). We thus established that the three known flavokavains (Figure 2B) are produced by the combinatorial activities of *Pm*CHS, *Pm*KOMT1, and *Pm*KOMT2.

**Figure 11.**
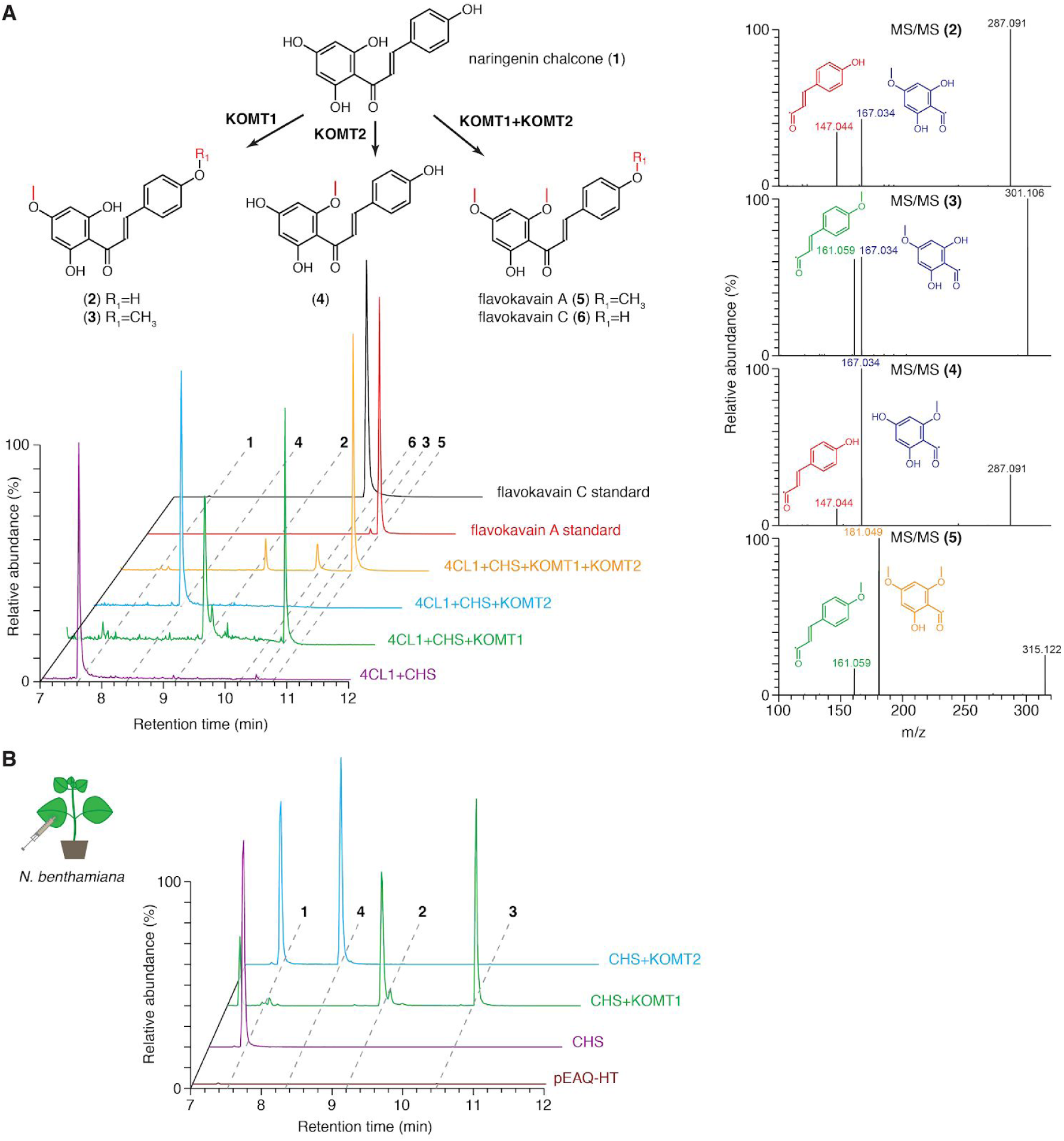
The biosynthesis of flavokavains using *Pm*CHS, *Pm*KOMT1, and *Pm*KOMT2. **A.** Combined LC-MS XICs of 273.076 m/z (**1**), 287.091 m/z (**2, 4**), 301.107 m/z (**3, 6**), and 315.123 m/z (**5**) from *in vitro* enzyme assays using the indicated combinations of enzymes and *p*-coumaric acid as the starter substrate. MS/MS analysis interpretation is shown on the right to support the identification of individual products**. B.** Combined LC-MS XICs of 273.076 m/z (**1**), 287.091 m/z (**2, 4**), 301.107 m/z (**3, 6**) in leaf extracts from Nicotiana transiently expressing indicated enzymes. Expression in Nicotiana resulted in equivalent compounds to those produced by *in vitro* enzyme assays.

As a proof of concept, we examined the feasibility of kavalactone production in microbial hosts through the means of metabolic engineering. Using an operon-like construct co-expressing *Pm4CL1* with *PmSPS1* or *PmSPS2* under the constitutive *pGAP* promoter, we could reconstitute the biosynthesis of the kavalactone precursor, bisnoryangonin, from supplemented *p*-coumaric acid in *E.coli* under shake flask condition (Figure 12A). Similarly, plasmid-based expression of the same combination of enzymes under the constitutive *pTEF* promoter in the yeast *Saccharomyces cerevisiae* also resulted in bisnoryangonin production from supplemented *p*-coumaric acid under shake flask condition (Figure 12B).

**Figure 12.**
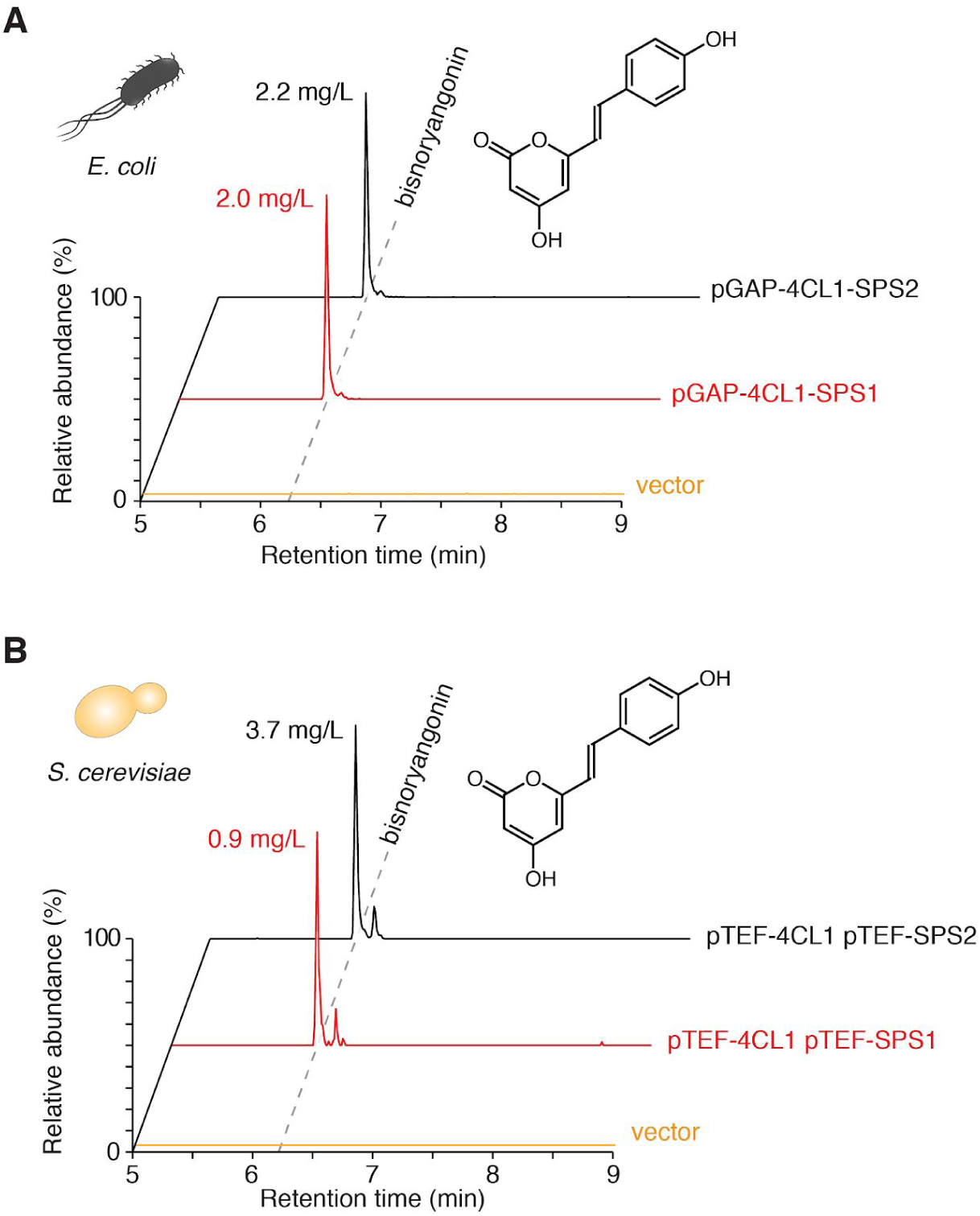
Heterologous production of the kavalactone precursor, bisnoryangonin, in *E. coli* and yeast. **A.** LC-MS XIC of 231.065 m/z (bisnoryangonin) from the media supernatants of *E. coli* cultures incubated overnight at 30°C with 1 mM *p*-coumaric acid. Cells were expressing the indicated enzymes from a single plasmid under the control of a constitutive *pGAP* promoter. Final bisnoryangonin yields are indicated next to each peak. **B.** LC-MS XIC of 231.065 m/z (bisnoryangonin) from the media supernatants of *S. cerevisiae* cultures incubated for 2 days at 30°C with 2 mM *p*-coumaric acid. Cells were expressing the indicated enzymes from two plasmids under the control of a constitutive *pTEF* promoter.

The kava extracts display complex pharmacology in animals, which include modulation of GABA_A_ and cannabinoid (CB_1_) receptors, blockade of voltage-gated sodium ion channels, reduced release and reuptake of several neurotransmitters, and interactions with monoamine oxidase B and liver cytochromes P450^4,5,26^. The capability to produce structurally diverse kavalactones and flavokavains using their biosynthetic enzymes described herein will help delineate the action mechanism and structure-function relationship of kavalactones. Furthermore, combinatorial biosynthesis using additional tailoring enzymes from other sources will further enable diversification of kavalactones for novel biomedical applications. In light of several major challenges facing modern society, such as anxiety, opioid crisis, and the scarcity of effective treatments for psychiatric disorders, kavalactones and their derivatives present a promising class of psychoactive molecules to potentially answer these unmet needs.

## Materials and Methods

### Chemicals and reagents

Kavalactone standards (yangonin, methysticin, desmethoxyyangonin, 7,8-dihydrokavain, and 7,8-dihydromethysticin) were obtained from AvaChem Scientific.

*p*-Coumaroyl-CoA was obtained from MicroCombiChem. Naringenin was obtained from AK Scientific. (±)-Kavain, malonyl-CoA, *trans*-cinnamic acid-d_6_, and other cofactors and reagents were obtained from Sigma-Aldrich.

### RNA extraction and cDNA template preparation

Total RNA was extracted separately from kava root and leaf tissue using the RNeasy Plant Mini Kit (QIAGEN). First-strand cDNAs were synthesized by RT–PCR from the total RNA samples as templates using the SuperScript III First-Strand Synthesis System with the oligo(dT)_20_ primer (Thermo Fisher Scientific).

### Transcriptome sequencing and assembly

The transcriptome library preparation and sequencing were performed at the Beijing Genomics Institute using the standard BGISEQ-500 RNA sample preparation protocol. The libraries were sequenced as 50×50 bp (PE-50) reads on the BGISEQ-500 platform. Sequence reads (FASTQ files) from leaf and root samples were merged into a single set of reads, trimmed for sequencing adaptors using Trimmomatic^29^, and assembled into a *de novo* transcriptome using Trinity^30^, resulting in 285,753 transcript sequences. Within those, we identified 302,366 putative open reading frames using Transdecoder^31^. We evaluated the transcriptome as 78.6% complete by the metric of Benchmarking Universal Single-Copy Orthologs (BUSCO)^32^. Gene expression statistics were determined using RSEM^33^. Transcripts and predicted protein sequences were annotated with transcript-per-million (TPM) values and closest BLAST hits from the UniProtKB/Swiss-Prot database using in-house scripts. Transcriptome mining was performed on a local BLAST server^34^. In addition, two kava root RNAseq datasets previously published by the PhytoMetaSyn Project^35^ (NCBI SRA accessions SRX202785 and SRX202184) were assembled and used as complementary sources of candidate enzyme sequences. Existing RNAseq datasets of *Piper auritum, Piper betle*, and *Piper nigrum* (NCBI SRA accessions ERX2099199, SRX691517, and SRX890122) were also assembled into *de novo* transcriptomes to obtain their corresponding CHS sequences.

### Sequence alignment and phylogenetic analyses

Sequence alignments were performed using the MUSCLE^36^ algorithm in MEGA7^37^. Evolutionary histories were inferred by using the Maximum Likelihood method based on the JTT matrix-based model^38^. Bootstrap values were calculated using 1,000 replicates. All phylogenetic analyses were conducted in MEGA7^37^.

### Evolutionary analysis

The Mixed Effects Model of Evolution (MEME) analysis^19^ was performed using the Hypothesis Testing using Phylogenies (HyPhy) package^39^, starting from codon-aligned CHS nucleotide sequences and the corresponding phylogenetic tree. The p-value threshold was set to 0.05.

### Cloning of candidate genes from cDNA

Phusion High-Fidelity DNA Polymerase (Thermo Fisher Scientific) was used for PCR amplifications from kava cDNA. Gibson assembly was used to clone the amplified genes into target vectors^40^. Restriction enzymes and Gibson assembly reagents were purchased from New England Biolabs. Oligonucleotide primers were purchased from Integrated DNA Technologies. All primers used for cloning are listed in Supplementary Table 3.

### Protein expression and purification

Candidate genes were cloned into pHis8-4, a bacterial expression vector containing an N-terminal 8×His tag followed by a tobacco etch virus (TEV) cleavage site for recombinant protein production in *E. coli*. Proteins were expressed in the BL21(DE3) *E. coli* strain cultivated in terrific broth (TB) and induced with 0.1 mM isopropyl β-D-1-thiogalactopyranoside (IPTG) overnight at 18 °C. For the COMTL enzyme screen, crude protein extracts were prepared from *E. coli* cultures using Bacterial Protein Extraction Reagent (B-PER, Thermo Fisher Scientific). For protein purification, *E. coli* cells were harvested by centrifugation, resuspended in 150 mL lysis buffer (50 mM Tris pH 8.0, 500 mM NaCl, 30 mM imidazole, 5 mM DTT), and lysed with five passes through an M-110L microfluidizer (Microfluidics). The resulting crude protein lysate was clarified by centrifugation (19,000 g, 1 h) prior to QIAGEN nickel–nitrilotriacetic acid (Ni–NTA) gravity flow chromatographic purification. After loading the clarified lysate, the Ni–NTA resin was washed with 20 column volumes of lysis buffer and eluted with 1 column volume of elution buffer (50 mM Tris pH 8.0, 500 mM NaCl, 300 mM imidazole, 5 mM DTT). 1 mg of His-tagged TEV protease^41^ was added to the eluted protein, followed by dialysis at 4 °C for 16 h in dialysis buffer (50 mM Tris pH 8.0, 500 mM NaCl, 5 mM DTT). After dialysis, protein solution was passed through Ni–NTA resin to remove uncleaved protein and His-tagged TEV. The recombinant proteins were further purified by gel filtration on a fast protein liquid chromatography (FPLC) system (GE Healthcare Life Sciences). The principal peaks were collected, verified by SDS–PAGE, and dialyzed into a storage buffer (12.5 mM Tris pH 8.0, 50 mM NaCl, 5 mM DTT). Finally, proteins were concentrated to >10 mg/mL using Amicon Ultra-15 Centrifugal Filters (Millipore). The *Pm*SPS1, *Pm*SPS2, *Pm*CHS, and *Pm*KLR1 proteins purified as homodimers, while *Pm*4CL1, *Pm*KOMT1 and *Pm*KOMT2 purified as monomers.

### *In vitro* enzyme assays

Enzyme assays were performed in 50 mM potassium phosphate buffer, pH 7.6 containing 5 mM MgCl_2_, 3 mM adenosine triphosphate (ATP), 1 mM coenzyme A, 3 mM malonyl-CoA, and 0.25 mM initial substrate (typically, cinnamic acid or *p*-coumaric acid).

Recombinant enzymes were added to final concentration of 10 µg/mL. For OMT assays, additional 3 mM S-adenosyl-methionine was added. For reductase assays, additional 5 mM reduced nicotinamide adenine dinucleotide phosphate (NADPH) was added. Reactions were incubated at 30 °C until completion (typically overnight), followed by the addition of methanol to 50% final concentration. Samples were centrifuged (13,000 g, 20 min) and supernatants were collected for LC-MS analyses.

### Transgenic Arabidopsis

Kava *PKS* genes were cloned into the pEarleyGate 100 vector^42^ and transformed into *Agrobacterium tumefaciens* strain GV3130. The *Arabidopsis thaliana tt4-2* mutant^17^ was then transformed using the *Agrobacterium*-mediated floral dip method^43^.

Transformants were selected by spraying with Finale (contains 11.33% glufosinate ammonium; Bayer) diluted 1:500 in water. Selection was repeated in each subsequent generation and experiments were performed in the T2 or T3 generations.

### qRT-PCR analysis

qRT-PCR reactions were performed on a QuantStudio 6 Flex system (Thermo Fisher Scientific) using SYBR Green Master Mix (Thermo Fisher Scientific) and primers listed in Supplementary Table 3. Gene expression Ct values were normalized using the reference gene *At1g13320^44^*.

### Plant metabolite extraction

Approximately 100 mg of plant leaf tissue was dissected, transferred into grinding tubes containing ~15 zirconia/silica disruption beads (2 mm diameter; Research Products International), and snap-frozen in liquid nitrogen. The frozen samples were homogenized twice on a TissueLyser II (QIAGEN). Metabolites were extracted using 5–10 volumes (w/v) of 50% methanol at 55 °C for 1 h. Extracts were centrifuged twice (13,000 g, 20 min) and supernatants were collected for LC-MS analysis.

### LC-MS analysis

LC was conducted on a Dionex UltiMate 3000 UHPLC system (Thermo Fisher Scientific), using water with 0.1% formic acid as solvent A and acetonitrile with 0.1% formic acid as solvent B. Reverse phase separation of analytes was performed on a Kinetex C18 column, 150 × 3 mm, 2.6 μm particle size (Phenomenex). Chiral chromatography was performed on a Lux i-Amylose-1 column, 250 × 4.6 mm, 5 µm particle size (Phenomenex). The column oven was held at 30 °C. Most injections were eluted with a gradient of 5–60% B for 9 min, 95% B for 3 min, and 5% B for 3 min, with a flow rate of 0.7 mL/min. The initial analysis of *Piper* species leaf extracts was performed on a gradient of 5–80% B for 40 min, 95% B for 4 min, and 5% B for 5 min, with a flow rate of 0.8 mL/min. Chiral chromatography was performed on a gradient of 5–95% B for 30 min, 95% B for 5 min, and 5% B for 10 min, with a flow rate of 1 mL/min. MS analyses for the OMT enzyme screen and for chiral chromatography were performed on a TSQ Quantum Access Max mass spectrometer (Thermo Fisher Scientific) operated in positive ionization mode with full scan range of 100–400 m/z. Other MS analyses and plant metabolic profiling were performed on a high-resolution Q-Exactive benchtop Orbitrap mass spectrometer (Thermo Fisher Scientific) operated in positive ionization mode with full scan range of 100–1000 m/z and top 5 data-dependent MS/MS scans. Raw LC-MS data were analyzed using XCalibur (Thermo Fisher Scientific), MZmine 2^45^, and MetaboAnalyst^46^. The SIRIUS tool was used to interpret MS/MS spectra^47,48^. Identified metabolites are listed in Supplementary Table 2.

### X-ray crystallography

The purified SPS1 protein (17.1 mg/mL) was incubated with 4 mM *p*-coumaroyl-CoA for 1 h prior to setting crystal trays. Crystals of *Pm*SPS1 were obtained after 4 days at 21 °C in hanging drops containing 0.8 μL of protein solution and 0.8 μL of reservoir solution (50 mM HEPES pH 7.5, 10% w/v PEG 8000, 4% v/v ethylene glycol). Crystals were frozen in reservoir solution with ethylene glycol concentration increased to 15% (v/v).

SPS1 X-ray diffraction data were collected on the 24-ID-C beam line equipped with a Pilatus-6MF pixel array detector (Advanced Photon Source, Argonne National Laboratory).

Crystals of the purified *Pm*CHS protein (17.6 mg/mL) were obtained from a screening plate after 4 days at 4°C in sitting drops containing 150 nL protein solution and 150 nL of reservoir solution (100 mM HEPES pH 7.0, 10% w/v PEG 6000). CHS X-ray diffraction data were collected on the 24-ID-E beam line equipped with an Eiger-16M pixel array detector (Advanced Photon Source, Argonne National Laboratory). Diffraction intensities were indexed and integrated with iMosflm^49^ and scaled with SCALA^50^. Initial phases were determined by molecular replacement using Phaser^51^ in PHENIX^52^, using search models generated from protein sequences on the Phyre^2^ server^53^. Subsequent structural building and refinements were conducted in PHENIX. Coot was used for graphical map inspection and manual tuning of atomic models^54^. Root mean square deviation (RMSD) of *Pm*SPS1 and *Pm*CHS structures was calculated using the Structure comparison function in PHENIX. Crystallography statistics are listed in Table 1.

### Protein structure modeling and rendering

The *Pm*SPS2 structure model was generated on the Phyre^2^ server using the *Pm*SPS1 X-ray structure as a modeling template^53^. The active site pocket in the *Pm*SPS1 structure was determined using KVFinder^55^. Molecular graphics were rendered with PyMOL (Schrödinger).

### Transient expression in *Nicotiana benthamiana*

Target kava genes were cloned into the pEAQ-HT vector^56,57^ and transformed into the ElectroMAX *Agrobacterium tumefaciens* strain LBA4404 (Invitrogen). Bacteria were cultivated at 30 °C to OD_600_ of 1.5 in 50 mL of YM medium (0.4 g/L yeast extract, 10 g/l mannitol, 0.1 g/L NaCl, 0.2 g/L MgSO_4·_7H_2_O, 0.5 g/L K_2_HPO_4·_3H_2_O), washed with 0.5×PBS buffer (68 mM NaCl, 1.4 mM KCl, 5 mM Na_2_HPO_4_, 0.9 mM KH_2_PO_4_), and resuspended in 0.5×PBS buffer to OD_600_ of 0.8. For co-expressing multiple genes, individual *A. tumefaciens* cultures containing the transgene constructs were grown, pelleted, and washed separately, and finally mixed to reach the final OD_600_ of 0.8 for each culture. 1 mL of the final culture was used to infiltrate the underside of 5-6 weeks old *N. benthamiana* leaves. Leaves were harvested 5 days post infiltration for metabolite extraction.

### Expression in *Escherichia coli*

*Pm4CL1*, *PmSPS1*, and *PmSPS2* were cloned into a custom-made, operon-like expression vector pJKW1565. The genes were preceded by a constitutively expressed *pGAP* promoter of the *E. coli* glyceraldehyde-3-phosphate dehydrogenase (GAPDH)^58^, and each gene was preceded by a 23-bp ribosome binding site (RBS) sequence^59^. *tSPY* was used as a terminator^60^. The constructed plasmids (pJKW1574 containing the *Pm4CL1* + *PmSPS1* operon and pJKW1658 containing the *Pm4CL1* + *PmSPS2* operon) were transformed into the BW27784 *E. coli* strain^61^ and cultivated at 30 °C overnight in TB medium supplied with 1 mM *p*-coumaric acid, followed by metabolite extraction from the culture fluid. An aliquot of the bacterial culture was mixed with an equal volume of methanol (50% final methanol concentration), centrifuged twice (13,000 g, 20 min) and supernatants were collected for LC-MS analysis. Since bisnoryangonin standard was not available, its concentrations were estimated using naringenin as the calibration-curve standard.

### Expression in *Saccharomyces cerevisiae*

The p42xTEF 2µ plasmids with various auxotrophic growth markers for constitutive expression in yeast were used^62^. The *Pm4CL1* gene was cloned into p425TEF (plasmid pJKW1413), while *PmSPS1* and *PmSPS2* were cloned into p426TEF (plasmids pJKW1538 and pJKW1547, respectively). The constructed plasmids were transformed into the BY4743 yeast strain^63^. Yeast cells were cultivated at 30 °C for 2 days in Yeast Nitrogen Base (YNB) + histidine + dextrose medium supplied with 2 mM *p*-coumaric acid, followed by metabolite extraction from the culture fluid. An aliquot of the yeast culture was mixed with an equal volume of methanol (50% final methanol concentration), centrifuged twice (13,000 g, 20 min) and supernatants were collected for LC-MS analysis.

## Acknowledgments

This work was supported by grants from the Edward N. and Della L. Thome Memorial Foundation, the Family Larsson-Rosenquist Foundation, and the National Science Foundation (CHE-1709616). T.P. is a Simons Foundation Fellow of the Helen Hay Whitney Foundation. J.K.W is supported by the Beckman Young Investigator Program, Pew Scholars Program in the Biomedical Sciences (grant number 27345), and the Searle Scholars Program (grant number 15-SSP-162). This work is based on research conducted at the Northeastern Collaborative Access Team (NE-CAT) beamlines, which are funded by the National Institute of General Medical Sciences from the National Institutes of Health (P41 GM103403). The Pilatus 6M detector on NE-CAT 24-ID-C beam line is funded by a NIH-ORIP HEI grant (S10 RR029205). This research used resources of the Advanced Photon Source, a U.S. Department of Energy (DOE) Office of Science User Facility operated for the DOE Office of Science by Argonne National Laboratory under Contract No. DE-AC02-06CH11357. RNAseq service was provided free of charge by the Beijing Genome Institute in exchange for an evaluation of their BGISEQ-500 sequencing platform. We thank Bryan Marotta for an introduction to kava, Chi Nguyen and Fadel A. Samatey for advice regarding crystallography, Gerald Fink for providing yeast strains and expression vectors, and Weng lab members for constructive comments. Kava plant photo courtesy of Randy Travis.

## Author Contributions

T.P. and J.K.W. designed experiments. T.P. performed experiments, while M.P.T.S. assisted with cloning and crystallography and T.R.F. assisted with transcriptome assembly and LC-MS analyses. A.D.A. cloned genes and purified proteins. C.H.S. constructed expression vectors. T.P. analysed data. T.P. and J.K.W. wrote the paper.

## References

1. Li, F.-S. & Weng, J.-K. Demystifying traditional herbal medicine with modern approach. Nature Plants 3, 17109 (2017).

2. Lebot, V. & Lévesque, J. The origin and distribution of kava (Piper methysticum Forst. f., Piperaceae): a phytochemical approach. Allertonia 5, 223–281 (1989).

3. Singh, Y. N. Kava: an overview. J. Ethnopharmacol. 37, 13–45 (1992).

4. Cairney, S., Maruff, P. & Clough, A. R. The neurobehavioural effects of kava. Aust. N. Z. J. Psychiatry 36, 657–662 (2002).

5. Sarris, J., LaPorte, E. & Schweitzer, I. Kava: a comprehensive review of efficacy, safety, and psychopharmacology. Aust. N. Z. J. Psychiatry 45, 27–35 (2011).

6. Chua, H. C. et al. Kavain, the Major Constituent of the Anxiolytic Kava Extract, Potentiates GABAA Receptors: Functional Characteristics and Molecular Mechanism. PLOS ONE 11, e0157700 (2016).

7. Jamieson, D. D. & Duffield, P. H. The antinociceptive actions of kava components in mice. Clin. Exp. Pharmacol. Physiol. 17, 495–507 (1990).

8. Depression and other common mental disorders: global health estimates. (Geneva: World Health Organization, 2017).

9. Zi, X. & Simoneau, A. R. Flavokawain A, a novel chalcone from kava extract, induces apoptosis in bladder cancer cells by involvement of Bax protein-dependent and mitochondria-dependent apoptotic pathway and suppresses tumor growth in mice. Cancer Res. 65, 3479–3486 (2005).

10. Zhou, P. et al. Flavokawain B, the hepatotoxic constituent from kava root, induces GSH-sensitive oxidative stress through modulation of IKK/NF-kappaB and MAPK signaling pathways. FASEB J. 24, 4722–4732 (2010).

11. Abe, I. & Morita, H. Structure and function of the chalcone synthase superfamily of plant type III polyketide synthases. Nat. Prod. Rep. 27, 809–838 (2010).

12. Lebot, V., Aradhya, M. K. & Manshardt, R. M. Geographic survey of genetic variation in kava (Piper methysticum Forst. f. and P. wichmannii C. DC.). Pacific Science 45, 169–185 (1991).

13. Torrens-Spence, M. P., Fallon, T. R. & Weng, J. K. A Workflow for Studying Specialized Metabolism in Nonmodel Eukaryotic Organisms. in Methods in Enzymology 69–97 (2016).

14. Owen, C., Patron, N. J., Huang, A. & Osbourn, A. Harnessing plant metabolic diversity. Curr. Opin. Chem. Biol. 40, 24–30 (2017).

15. Tatsis, E. C. & O’Connor, S. E. New developments in engineering plant metabolic pathways. Curr. Opin. Biotechnol. 42, 126–132 (2016).

16. Akiyama, T., Shibuya, M., Liu, H. M. & Ebizuka, Y. p-Coumaroyltriacetic acid synthase, a new homologue of chalcone synthase, from Hydrangea macrophylla var. thunbergii. FEBS J. 263, 834–839 (1999).

17. Burbulis, I. E., Iacobucci, M. & Shirley, B. W. A null mutation in the first enzyme of flavonoid biosynthesis does not affect male fertility in Arabidopsis. Plant Cell 8, 1013–1025 (1996).

18. Sainsbury, F. & Lomonossoff, G. P. Transient expressions of synthetic biology in plants. Curr. Opin. Plant Biol. 19, 1–7 (2014).

19. Murrell, B. et al. Detecting individual sites subject to episodic diversifying selection. PLoS Genet. 8, e1002764 (2012).

20. Jez, J. M. et al. Structural control of polyketide formation in plant-specific polyketide synthases. Chem. Biol. 7, 919–930 (2000).

21. Morita, H. et al. A structure-based mechanism for benzalacetone synthase from Rheum palmatum. Proc. Natl. Acad. Sci. U. S. A. 107, 669–673 (2010).

22. Austin, M. B. & Noel, J. P. The chalcone synthase superfamily of type III polyketide synthases. Nat. Prod. Rep. 20, 79–110 (2003).

23. Moinuddin, S. G. A. et al. Insights into lignin primary structure and deconstruction from Arabidopsis thaliana COMT (caffeic acid O-methyl transferase) mutant Atomt1. Org. Biomol. Chem. 8, 3928–3946 (2010).

24. Eckermann, C. et al. Stilbenecarboxylate biosynthesis: a new function in the family of chalcone synthase-related proteins. Phytochemistry 62, 271–286 (2003).

25. Gosch, C., Halbwirth, H. & Stich, K. Phloridzin: biosynthesis, distribution and physiological relevance in plants. Phytochemistry 71, 838–843 (2010).

26. Li, Y. et al. Methysticin and 7,8-dihydromethysticin are two major kavalactones in kava extract to induce CYP1A1. Toxicol. Sci. 124, 388–399 (2011).

27. Ikezawa, N. et al. Molecular cloning and characterization of CYP719, a methylenedioxy bridge-forming enzyme that belongs to a novel P450 family, from cultured Coptis japonica cells. J. Biol. Chem. 278, 38557–38565 (2003).

28. Lau, W. & Sattely, E. S. Six enzymes from mayapple that complete the biosynthetic pathway to the etoposide aglycone. Science 349, 1224–1228 (2015).

29. Bolger, A. M., Lohse, M. & Usadel, B. Trimmomatic: a flexible trimmer for Illumina sequence data. Bioinformatics 30, 2114–2120 (2014).

30. Grabherr, M. G. et al. Full-length transcriptome assembly from RNA-Seq data without a reference genome. Nat. Biotechnol. 29, 644–652 (2011).

31. Haas, B. J. et al. De novo transcript sequence reconstruction from RNA-seq using the Trinity platform for reference generation and analysis. Nat. Protoc. 8, 1494–1512 (2013).

32. Simão, F. A., Waterhouse, R. M., Ioannidis, P., Kriventseva, E. V. & Zdobnov, E. M. BUSCO: assessing genome assembly and annotation completeness with single-copy orthologs. Bioinformatics 31, 3210–3212 (2015).

33. Li, B. & Dewey, C. N. RSEM: accurate transcript quantification from RNA-Seq data with or without a reference genome. BMC Bioinformatics 12, 323 (2011).

34. Priyam, A. et al. Sequenceserver: a modern graphical user interface for custom BLAST databases. bioRxiv 033142 (2015). doi:10.1101/033142

35. Xiao, M. et al. Transcriptome analysis based on next-generation sequencing of non-model plants producing specialized metabolites of biotechnological interest. J. Biotechnol. 166, 122–134 (2013).

36. Edgar, R. C. MUSCLE: multiple sequence alignment with high accuracy and high throughput. Nucleic Acids Res. 32, 1792–1797 (2004).

37. Kumar, S., Stecher, G. & Tamura, K. MEGA7: Molecular Evolutionary Genetics Analysis Version 7.0 for Bigger Datasets. Mol. Biol. Evol. 33, 1870–1874 (2016).

38. Jones, D. T., Taylor, W. R. & Thornton, J. M. The rapid generation of mutation data matrices from protein sequences. Comput. Appl. Biosci. 8, 275–282 (1992).

39. Pond, S. L. K., Frost, S. D. W. & Muse, S. V. HyPhy: hypothesis testing using phylogenies. Bioinformatics 21, 676–679 (2005).

40. Gibson, D. G. et al. Enzymatic assembly of DNA molecules up to several hundred kilobases. Nat. Methods 6, 343–345 (2009).

41. Tropea, J. E., Cherry, S. & Waugh, D. S. Expression and purification of soluble His(6)-tagged TEV protease. Methods Mol. Biol. 498, 297–307 (2009).

42. Earley, K. W. et al. Gateway-compatible vectors for plant functional genomics and proteomics. Plant J. 45, 616–629 (2006).

43. Clough, S. J. & Bent, A. F. Floral dip: a simplified method forAgrobacterium-mediated transformation ofArabidopsis thaliana: Floral dip transformation of Arabidopsis. Plant J. 16, 735–743 (1998).

44. Czechowski, T., Stitt, M., Altmann, T., Udvardi, M. K. & Scheible, W.-R. Genome-wide identification and testing of superior reference genes for transcript normalization in Arabidopsis. Plant Physiol. 139, 5–17 (2005).

45. Pluskal, T., Castillo, S., Villar-Briones, A. & Oresic, M. MZmine 2: modular framework for processing, visualizing, and analyzing mass spectrometry-based molecular profile data. BMC Bioinformatics 11, 395 (2010).

46. Xia, J., Sinelnikov, I. V., Han, B. & Wishart, D. S. MetaboAnalyst 3.0–making metabolomics more meaningful. Nucleic Acids Res. 43, W251–W257 (2015).

47. Böcker, S., Letzel, M. C., Lipták, Z. & Pervukhin, A. SIRIUS: decomposing isotope patterns for metabolite identification. Bioinformatics 25, 218–224 (2009).

48. Dührkop, K., Shen, H., Meusel, M., Rousu, J. & Böcker, S. Searching molecular structure databases with tandem mass spectra using CSI:FingerID. Proc. Natl. Acad. Sci. U. S. A. 112, 12580–12585 (2015).

49. Battye, T. G. G., Kontogiannis, L., Johnson, O., Powell, H. R. & Leslie, A. G. W. \it iMOSFLM: a new graphical interface for diffraction-image processing with \it MOSFLM. Acta Crystallogr. D Biol. Crystallogr. 67, 271–281 (2011).

50. Evans, P. Scaling and assessment of data quality. Acta Crystallogr. D Biol. Crystallogr. 62, 72–82 (2006).

51. McCoy, A. J. Solving structures of protein complexes by molecular replacement with Phaser. Acta Crystallogr. D Biol. Crystallogr. 63, 32–41 (2007).

52. Adams, P. D. et al. PHENIX: a comprehensive Python-based system for macromolecular structure solution. Acta Crystallogr. D Biol. Crystallogr. 66, 213–221 (2010).

53. Kelley, L. A., Mezulis, S., Yates, C. M., Wass, M. N. & Sternberg, M. J. E. The Phyre2 web portal for protein modeling, prediction and analysis. Nat. Protoc. 10, 845–858 (2015).

54. Emsley, P. & Cowtan, K. Coot: model-building tools for molecular graphics. Acta Crystallogr. D Biol. Crystallogr. 60, 2126–2132 (2004).

55. Oliveira, S. H. P. et al. KVFinder: steered identification of protein cavities as a PyMOL plugin. BMC Bioinformatics 15, 197 (2014).

56. Sainsbury, F., Thuenemann, E. C. & Lomonossoff, G. P. pEAQ: versatile expression vectors for easy and quick transient expression of heterologous proteins in plants. Plant Biotechnol. J. 7, 682–693 (2009).

57. Peyret, H. & Lomonossoff, G. P. The pEAQ vector series: the easy and quick way to produce recombinant proteins in plants. Plant Mol. Biol. 83, 51–58 (2013).

58. Charpentier, B., Bardey, V., Robas, N. & Branlant, C. The EIIGlc protein is involved in glucose-mediated activation of Escherichia coli gapA and gapB-pgk transcription. J. Bacteriol. 180, 6476–6483 (1998).

59. Olins, P. O. & Rangwala, S. H. A novel sequence element derived from bacteriophage T7 mRNA acts as an enhancer of translation of the lacZ gene in Escherichia coli. J. Biol. Chem. 264, 16973–16976 (1989).

60. Chen, Y.-J. et al. Characterization of 582 natural and synthetic terminators and quantification of their design constraints. Nat. Methods 10, 659–664 (2013).

61. Khlebnikov, A., Datsenko, K. A., Skaug, T., Wanner, B. L. & Keasling, J. D. Homogeneous expression of the P(BAD) promoter in Escherichia coli by constitutive expression of the low-affinity high-capacity AraE transporter. Microbiology 147, 3241–3247 (2001).

62. Mumberg, D., Müller, R. & Funk, M. Yeast vectors for the controlled expression of heterologous proteins in different genetic backgrounds. Gene 156, 119–122 (1995).

63. Brachmann, C. B. et al. Designer deletion strains derived from Saccharomyces cerevisiae S288C: a useful set of strains and plasmids for PCR-mediated gene disruption and other applications. Yeast 14, 115–132 (1998).

64. Ferrer, J. L., Jez, J. M., Bowman, M. E., Dixon, R. A. & Noel, J. P. Structure of chalcone synthase and the molecular basis of plant polyketide biosynthesis. Nat. Struct. Biol. 6, 775–784 (1999).

65. Veit, M. & Pauli, G. F. Major flavonoids from Arabidopsis thaliana leaves. J. Nat. Prod. 62, 1301–1303 (1999).

